# Btbd6-dependent Plzf recruitment to Cul3 E3 ligase complexes through BTB domain heterodimerization

**DOI:** 10.1101/575910

**Authors:** Mohamed Ismail, Stephen R. Martin, Neil J. Ball, Ian A. Taylor, Steven Howell, David G. Wilkinson, Stephen J. Smerdon

**Author notes:** **Corresponding Author**: Stephen Smerdon, **Corresponding Author’s Institution**: Francis Crick Institute.

## Abstract

The Cul3 adaptor Btbd6 plays crucial roles in neural development by driving the ubiquitin-dependent degradation of promyelocytic zinc finger transcription factor (Plzf). Btbd6 has conserved motifs, BTB-BACK-PHR, and by analogy with other BTB-BACK adaptors, might be expected to bind to Cul3 through the BTB-BACK domain, and to substrate through the PHR domain. However, we now present a mode of adaptor-substrate interaction through heterodimerisation between the normally homodimeric BTB domains of Btbd6 and Plzf. This heterodimerization appears to occur through monomer exchange that is detected only at or near physiological concentrations. The Btbd6-Plzf heterodimer thus formed assembles into a ternary complex with Cul3. In addition we show that the BTB and PHR domains of Btbd6 promote localisation in the nucleus and that the BACK domain contains a nuclear export signal. Our findings support a model whereby Btbd6 moves into and out of the nucleus, iteratively ‘sweeping’ Plzf into the cytoplasm and enabling complex formation with Cul3 that presents Plzf for ubiquitination.

**Highlights:**

- A general mechanism for recruitment of BTB domain-containing substrates by BTBdomain adaptors for the Cul3 E3 ligase complex
- Nuclear export of the Plzf/Btbd6 complex mediated by a NES within the Btbd6 BACK domain
- Cul3-dependent Plzf ubiquitylation through heterodimerisation of BTB domains on adaptor and substrate by monomer exchang

## Introduction

The targeted degradation of specific proteins is a key regulatory process in many biological signaling systems. This is mediated by the K48-linked polyubiquitination of proteins that marks them for degradation in the proteasome [1, 2]. In the largest family of E3 ligases, a cullin acts as a scaffold for the RING finger gene Rbx1 which recruits the E2 enzyme, and for a linker/adaptor complex that mediates binding to the substrate protein [3]. The cullins Cul1, Cul2 and Cul3 share the same overall architecture and ubiquitination mechanisms but differ in their substrate recognition modules [4–6]. In the case of Cul1, Skp1 is the linker between the cullin and an F box adaptor protein that binds the substrate [7]. The Cul2 complex has a similar structure, in which Elongin C is the linker and a SOCS box protein the adaptor [8]. In the case of Cul3, the adaptor proteins have a BTB domain and adjacent BACK domain which mediate direct binding to the cullin [8, 9].

The Cul3 adaptors studied in most detail recognize their substrate through a MATH domain or Kelch repeats [6, 10, 11]. Studies of these adaptors have found that a socalled ‘3-box’ motif in the BACK domain of SPOP and other BTB-BACK containing proteins greatly increases the interaction of the BTB domain with Cul3 [9, 12]. The crystal structure of KLHL11-Cul3 complex revealed the contribution of the 3-box motif to the interaction with Cul3 [13]. Indeed, it has been suggested that BTB proteins lacking a 3-box motif may not function as Cul3 adaptors [9]. In the case of SPOP, the BACK domain is also involved in high order oligomerization [9, 12], but it is not known whether this is the case for BTB-BACK-Kelch type adaptors [14].

A third family of BTB-BACK proteins likely to act as Cul3 adaptors have a PHR domain instead of the Kelch repeats or MATH domain [15–17]. Vertebrate genomes contain four genes of this BTB-BACK-PHR family: Btbd1-3 and Btbd6. Several studies have found potential substrates for members of this family [15, 18, 19], but little is known regarding the function or structural basis of the interactions. In our previous work, a yeast two-hybrid screen for proteins that bind Btbd6 identified both Cul3 and a transcriptional repressor protein, Plzf [15]. Plzf is a member of the BTB ZF family of proteins, and contains an N-terminal BTB domain and nine C-terminal Cys2-His2 Krüppel zinc fingers [17, 20, 21]. Immunoprecipitation studies showed that Btbd6 binding to Cul3 and Plzf leads to ubiquitination and degradation of Plzf, promoting neurogenesis by relieving Plzf-mediated inhibition of neuronal differentiation [15]. In addition, Btbd6 was found to promote nuclear export of Plzf and thus block its activity, analogous to previous findings for the adaptor protein KEAP1 and Nrf2 transcription factor [22].

Together, these observations reveal a mechanism in which Btbd6 enters the nucleus, binds to Plzf and promotes its export to the cytoplasm where, as a complex with Cul3, it is targeted for degradation. By analogy with other BTB-BACK adaptors, it might be assumed that the BTB-BACK of Btbd6 binds to Cul3 while the PHR domain binds to substrate. However, immunoprecipitation analysis of deletion mutants of Btbd6 unexpectedly suggests that the BTB domain and not the PHR domain of the adaptor is required for binding to Plzf [15]. This observation raises the possibility that there is a distinct mode of adaptor activity specifically tailored to BTB domain-containing substrates such as Plzf. Here we use a combination of biochemical, biophysical and cellbiological approaches to reveal a previously unseen mode of adaptor function in recruitment of BTB domain-containing substrates to cullin-dependent E3 ligase complexes.

## Results

### Plzf interacts with the Btbd6-Cul3 complex in vitro

In order to investigate assembly of the Btbd6-Plzf-Cul3 complex we set out to determine an interaction map for the three components using a panel of protein domains (Figure 1A) in binding affinity measurements employing pull-down assays, isothermal titration calorimetry (ITC) and microscale thermophoresis (MST). Studies of other adaptors have found that the BTB domain and 3-box alpha helical motif in the BACK domain mediate binding to Cul3 [9]. Indeed, the 32 amino acids immediately C-terminal to the Btbd6 BTB domain and within the BACK domain showed significant sequence similarities with the 3-box motif in other Cul3 adaptors (Figure 1B). Consistent with this idea, we observed a ~100-fold increase in affinity for binding of a BTB-BACK fragment of Btbd6 to the N-terminal region of Cul3 (Cul3-NTD: residues 1 −379) compared with the BTB domain alone (Figure 1C, S1 A & B). We also obtained a similar result on complex formation using Size Exclusion Chromatography Multi-Angle Light Scattering (SEC-MALS) (S2 A & B). Furthermore, and consistent with findings for other Cul3 adaptors [13], we found that Cul3 NTD lacking the first 18 amino acids (Cul3 NTD Δ18) has a dramatically reduced affinity for the BTB-BACK of Btbd6 (Figure 1D, S1 C and D). In contrast, the PHR domain of Btbd6 showed no interaction with Cul3 as evidenced by ITC, MST or protein pull-down assays (data not shown), suggesting that the interaction of Cul3 with the Btbd6 adaptor is entirely mediated by the BTB-BACK region.

**Figure 1.**
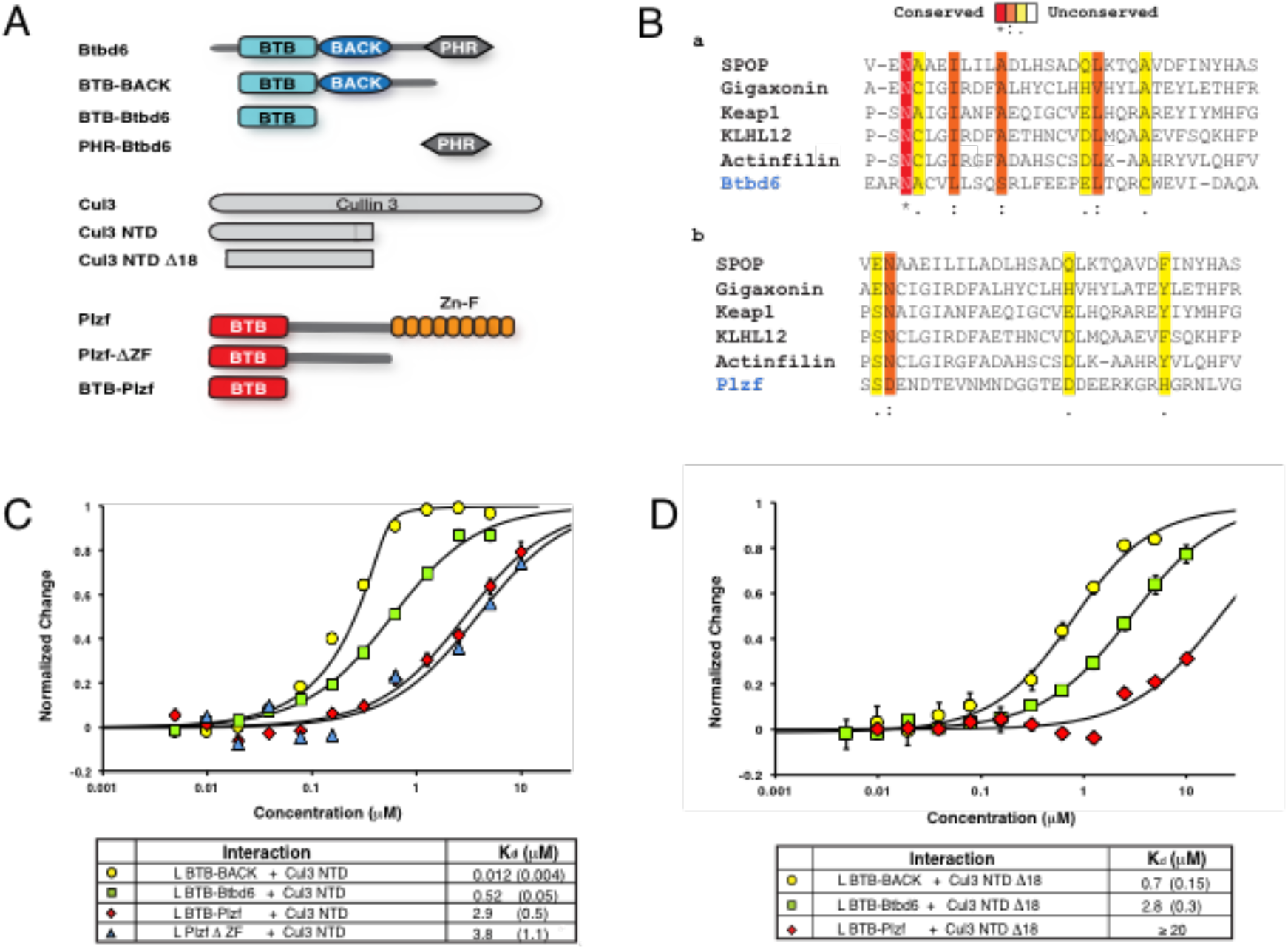
Interaction domain mapping. A Schematic showing the domain structures of Btbd6, Plzf and Cul3 and the deletions that have been used in this study. B Sequence alignment of the 32 residues following the BTB domain of Btbd6 (a) and Plzf (b) with other Cul3 adaptor proteins. Btbd6 shows strong sequence similarity with other adaptors in the so-called 3-box region, whereas Plzf showed no significant similarity. C MST measurements using labeled (designated as ‘L’) BTB BACK, BTB-Btbd6, BTB-Plzf and Plzf ΔZF with unlabeled Cul3 NTD. D MST data showing that unlabeled Cul3 NTD Δ18 binds more weakly to all three proteins L BTB-Btbd6, L BTB-BACK and L BTB-Plzf. E MST analysis of ternary complex formation.

The BTB domain of Plzf has been suggested to interact directly with Cul3 [23, 24], as well as indirectly via Btbd6 [15]. In order to address which of these interactions is more physiologically relevant, we first expressed and purified the BTB domain of Plzf (BTBPlzf) and measured the affinity of interaction with Cul3 NTD. Quantitative analysis by MST showed that BTB-Plzf interacts with Cul3 NTD with a Kd of 2.9 μM which is around 6-fold weaker than the interaction of BTB-Btbd6 with Cul3, but, most significantly, some 240-fold weaker than that for the BTB-BACK of Btbd6 (Figure 1C). In contrast to Btbd6, addition of adjacent sequences from the linker region of Plzf had no significant effect on binding (Figure 1A). Presumably reflecting the lack of a 3-box-like motif, and the absence of alpha helical structure when analyised using circular dichroism (Figure 1B & S2 C and D). As seen for Btbd6, the Cul3 NTD Δ18 deletion interacted more weakly with BTB-Plzf, with a Kd ≥ 20 μM (Figure 1D). From these results, we conclude that Plzf can interact directly with Cul3 in a manner that is structurally similar to the Cul3 interaction with the Btbd6 BTB domain [12, 13]. However, the observation that this interaction is two orders of magnitude weaker than that observed for the Btbd6 BTB-BACK combination suggests that direct binding of Plzf by Cul3 is unlikely to play a significant role in Cul3-mediated Plzf degradation.

### Btbd6 and Plzf directly interacts through their BTB domains

The foregoing data suggest that Cul3-mediated Plzf degradation proceeds not through direct recruitment of Plzf but through its interaction with the Btbd6 adaptor. Therefore, we next investigated which domains of Btbd6 and Plzf mediate binding. Previous coimmunoprecipitation studies indicated that Btbd6 binding to Plzf requires the Btbd6 BTB domain [15]. These observations were confirmed and extended using MST to study the interaction of labeled BTB-Plzf with BTB-Btbd6 and BTB-BACK. The MST binding isotherm showed two phases when labeled BTB-Plzf was titrated with BTBBACK over a concentration range of 120 nM - 250 μM (Figure 2A). Curve fitting using only the low concentration data (120 nM-15 μM) showed a strong interaction with an apparent Kd of ~470 nM (Figure 2A) whilst analysis using only the high concentration range (2 μM - 250 μM) showed a very much weaker interaction (Kd > 500 μM). This suggests that the BTB domains of Plzf and Btbd6 may also oligomerise through low affinity interactions that are likely of no physiological consequence. In order to confirm the tight interaction between the two BTB domains, we repeated the MST experiment using labeled BTB-Plzf and titrating with BTB-BACK at low concentrations ranging from 5 nM - 10 μM. These data show that the two proteins do indeed interact strongly with a similar apparent affinity as observed in the previous experiment (Kd ~ 1 μM) (Figure 2B). A similar affinity was obtained when using labeled BTB-Plzf with BTBBtbd6 (Kd ~ 1.3 μM), suggesting that the Btbd6 BACK domain is not involved in this interaction. Importantly, we also observed the same interaction in the inverse MST titration of BTB-Plzf using labeled BTB-Btbd6 (Figure 2C).

**Figure 2.**
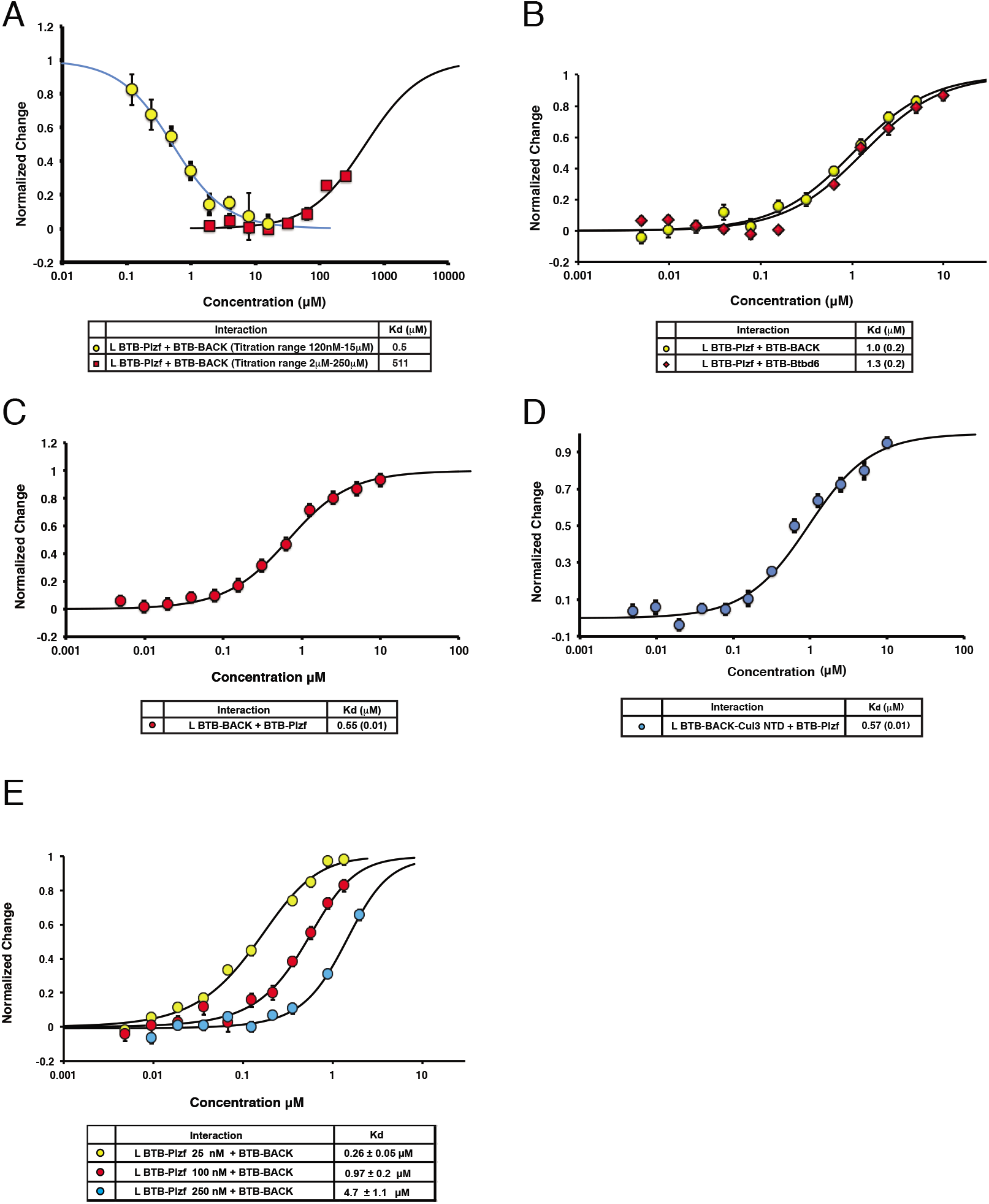
BTB-BTB interactions. A The interaction of L-BTB-Plzf with BTB shows two distinct phases of interaction. The first phase occurs at low concentrations of added BTB (120 nM-15 μM) with a Kd of 470 nM, and the second phase occurs at higher concentrations (15 μM – 250 μM) with a much weaker affinity (Kd ~ 500 μM). B Titrations of L-BTB-Plzf with BTBBtbd6 (red) and BTB-BACK (yellow). C Titration of L-BTB-BACK with BTB-Plzf.

The Cul3 NTD has been previously shown to interact with a surface on BTB domains that is distinct from that involved in BTB homodimerization [9]. Since the interaction between BTB-BACK and Cul3 NTD occurs with high affinity (Kd 12 nM), it was possible to use it as a stable complex to study the interaction with Plzf. BTB-BACK and Cul3 NTD were mixed together at a 1:1 ratio and the stoichiometric complex purified by gel filtration and labeled for MST analysis. The MST data show that labeled BTBBACK-Cul3 NTD complex at a concentration of 240 nM indeed interacts with BTB-Plzf with an apparent equilibrium dissociation constant of 570 nM (Figure 2D). Thus, BTBBtbd6 can simultaneously interact with Cul3 NTD and BTB-Plzf through two distinct and non-overlapping interfaces to form the physiologically relevant Cul3/Btbd6/Plzf complex.

In order to test this hypothesis we performed MST experiments with labeled BTB-Plzf at three different concentrations: 250, 100, and 25 nM (where BTB-Plzf will be approximately 22, 32, and 53% monomeric, respectively). Each was titrated with BTBBtbd6 concentrations ranging from 5 nM-10 μM. The apparent Kd for the interaction between BTB-Plzf and BTB-Btbd6 increased from ~260 nM (with 25 nM BTB-Plzf) to ~1 μM (with 100 nM BTB-Plzf) and ~4.7 μM (with 250 nm BTB-Plzf) (Figure 3 c). Such behavior is explicable by interaction of BTB-Plzf with monomeric but not dimeric BTB-Btbd6 to form a heterodimer (Figure 2E).

**Figure 3.**
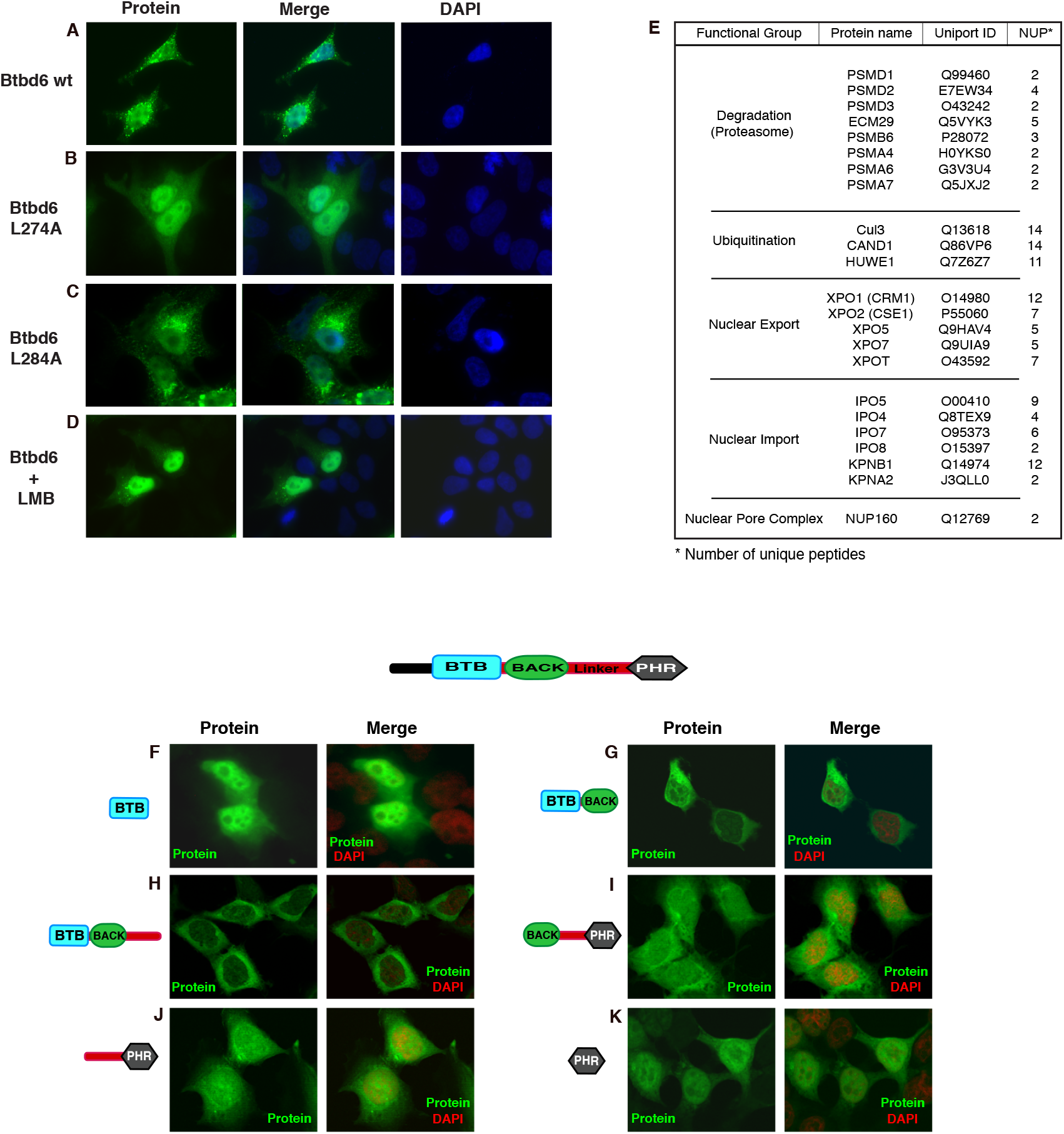
Nuclear/cytoplasmic shuttling by Btbd6. A (a) Btbd6wt is enriched in the cytoplasm and concentrated around the nuclear membrane. (b) Btbd6 274 L/A mutation results in a diffuse staining pattern and predominantly nuclear Btbd6 localization, in contrast to the punctate phenotype observed for Btbd6wt (c) Btbd6 284 L/A had no significant effect on Btbd6 localization. (d) LMB treatment retains Btbd6 in the nucleus, suggesting that Btbd6 LR-NES is dependent on CRM1 activity. B Mass spectrometry analysis of Btbd6 interactors. The most commonly observed co-immunoprecipitated proteins are classified into functional groups. NUP presents number of unique peptides. C Different deletions of Btbd6 were cloned in pEGFP1 that adds a GFP at the C-terminus of the expressed protein. The BTBBACK-Linker construct (h) shows a stronger cytoplasmic localization than BTB-BACK (g) locating NES activity to the BTB-BACK linker.

### Different domains of Btbd6 enable nuclear import and export

Btbd6 was found to promote Plzf relocalisation from the nucleus to the cytoplasm in zebrafish embryos [15] suggesting that Btbd6 might contain a nuclear export signal (NES). To test this possibility we used the NetNES program that searches for leucinerich NES (LR-NES) based on the sequence consensus L-x-(2,3)-[LIVFM]-x(2,3)-L-x- [LI] together with other characteristic properties [25, 26]. The NetNES sequence analysis of Btbd6 predicted a LR-NES motif in the BACK domain from L274 to L284 (170-AELALRSEGFSEIDLPTLE −288).

To address if L274 and L284 are indeed part of an NES motif, we made alanine substitutions of each of the two residues in full-length Btbd6 and expressed the three proteins (Btbd6-wt, Btbd6-L274A and Btbd6-L284A) as GFP fusions in HEK293 cells to visualise their subcellular localization. Wild-type Btbd6 was localised in a punctate subcellular distribution enriched in the cytoplasm and around the nuclear membrane, similar to the Cul3 adaptor KLHL7 [27] (Figure 3A a). Although Btbd6 L284A showed only a minor change in the subcellular distribution, the L274A mutant was mainly localised in the nucleus (Figure 3A b & c), identifying L274 as crucial for the function of the NES motif. Since export mediated by a leucine-rich NES depends upon CRM1 [28–32], we tested the effect of treating HEK293 cells over-expressing wild-type Btbd6 with a CRM1-specific export inhibitor leptomycin B (LMB). Indeed, after 3 hours of LMB treatment we observed that Btbd6 was mainly nuclear (Figure 3A d). Thus, Btbd6 contains a leucine-rich NES within the BACK domain that mediates CRM1-dependent nuclear export. To directly address whether Btbd6 undergoes nuclear import and export, myc-Btbd6 was expressed and immunoprecipitated from HEK293 cells, followed by mass spectrometry analysis. Among the interactors identified are proteins involved in nuclear export (CRM1 and CSE1) and import (IPO5 and KPNB1) and the nuclear pore complex protein NUP160 (Figure 3B).

To determine whether other sequences are present that regulate subcellular distribution, we generated fusions between individual domains of Btbd6 and GFP. We found that the BTB domain fusion protein was predominantly localised in the nucleus, whereas the BTB-BACK and BTB-BACK-linker fusion proteins were largely cytoplasmic, similar to the distribution of full length Btbd6 (Figure 3C). This finding is consistent with the BTB domain promoting nuclear import and the presence of an NES in the BACK domain. Likewise, the PHR and linker-PHR fusions were mainly found in the nucleus, while the subcellular distribution of the BTB-linker-PHR fusion was shifted towards the cytoplasm. However, the cytoplasmic versus nuclear distribution was much greater for BTB-BACK than for BACK-linker-PHR. This may reflect a different balance of nuclear import and export for these different combinations of domains, and/or that the ability of BTB-BACK to bind strongly to Cul3 in the cytoplasm contributes to the steady state subcellular distribution. Taken together, these results provide evidence that Btbd6 is a shuttle protein in which the BTB domain and PHR domain each promote nuclear import and the BACK domain contains a nuclear export signal.

## Discussion

Specific targeting of proteins for ubiquitin-dependent degradation is largely achieved through the recognition of substrates by a large family of E3 ubiquitin ligases that number over 200 in human cells [5]. These can be classified into distinct sub-families of structurally related proteins and complexes that recruit their target proteins in rather different ways and in response to a variety of different signals. We have previously shown that nuclear levels of the transcription repressor Plzf are down-regulated by export and Ub-dependent degradation by the Cul3 ubiquitin ligase adaptor Btbd6 leading directly to primary neural differentiation in zebrafish [15]. We now show that Btbd6-dependent degradation of Plzf occurs through a Cul3 adaptor-substrate recognition mechanism that seems to be specifically tailored to recognition of BTB containing substrates.

In our efforts to more precisely define the role of each of the structural domains within Btbd6 and Plzf, we were unable to detect any significant association of Btbd6 and Plzf by ITC. However, we were able to discern two forms of interaction using microscale thermophoresis comprising a low affinity binding of Btbd6 and Plzf BTB-domain dimers, and a second high-affinity interaction observed in a much lower concentration range that appears to be consistent with protomer exchange between Btbd6 and Plzf BTB domain dimers to form heterodimeric complexes. Indeed recent structural analyses of covalently linked BTB pairs shows how conservation of residues at the homodimer interface in a subset of BTB-domain family members can accommodate protomer exchange [33] consistent with our observation of functionally significant Btbd6/Plzf BTB domain heterodimerisation (Figure 4A).

**Figure 4.**
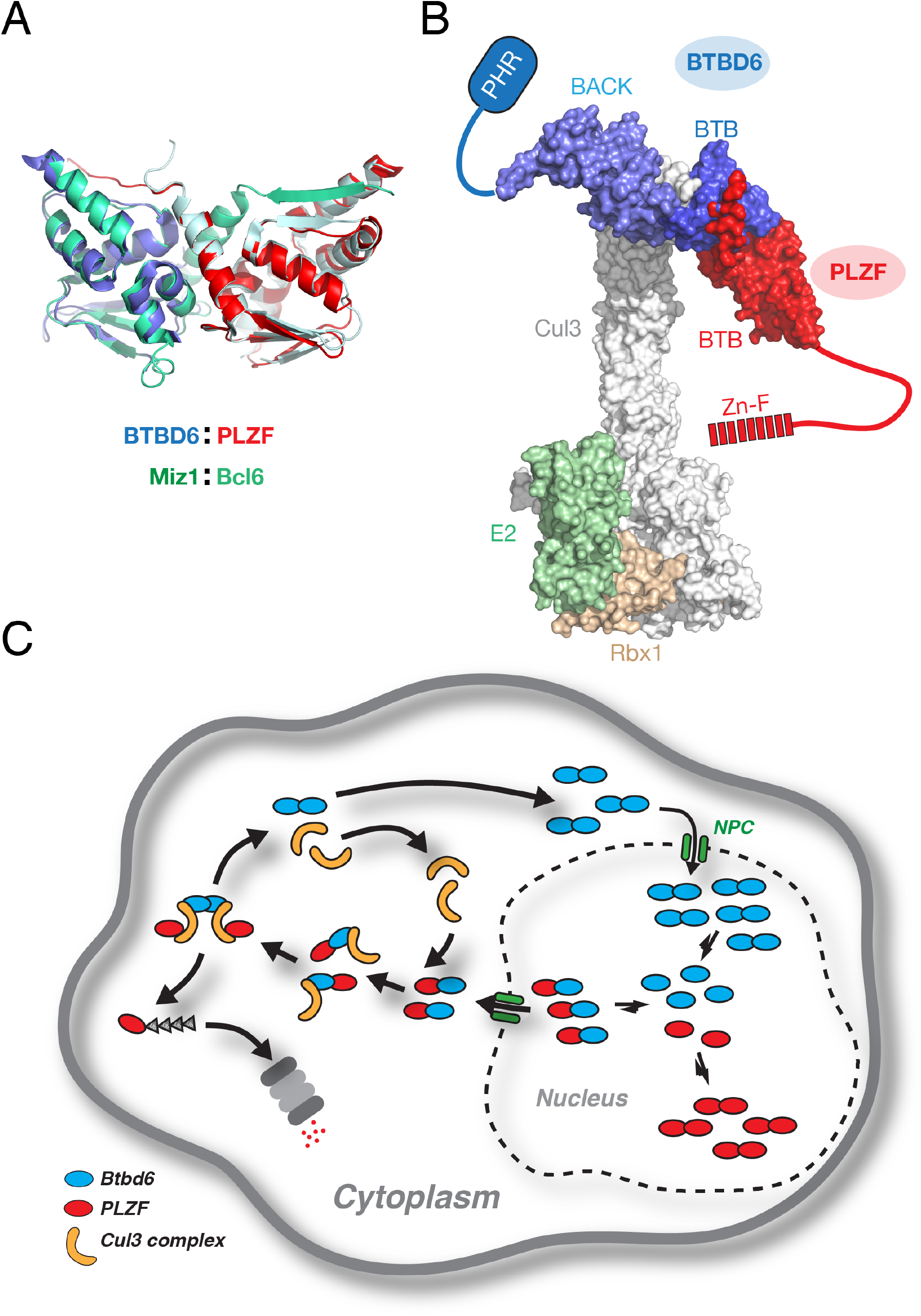
Pathway for Btbd6-mediated Plzf degradation. A Structural model of the Btbd6-Plzf BTB domain heterodimer. Btbd6 and Plzf monomer structures (PDB:2VKP & 1BUO respectively) were superimposed on the crystal structure of a tethered Miz1-Bcl6 heterodimer (PDB:4U2M) [33]. B The structural comparisons shown in panel A allows the generation of a model of the Plzf bound Cul3 complex C Schematic summary of the iterative removal of Plzf from the nuclear compartment through heterodimerisation with Btbd6, association with the Cul3 apparatus, Plzf ubiquitylation and degradation, and re-entry of Btbd6 into the nucleus.

Since the primary interaction site of the Cul3 N-terminal region is with a distinct surface formed between the BTB and BACK domains [9], Btbd6/Plzf heterodimers can form in the presence of Cul3 defining the heterotrimeric complex (Figure 2D) that appears to be the species active in Plzf ubiquitination. These observations, along with available structural data suggest a plausible molecular model of the substrate-bound Cul3/E2 complex (Figure 4B). In this assembly, the PHR domain extends away from the core cullin/adaptor complex, consistent with the lack of PHR binding to Cul3, while the Plzf substrate is recruited through the Btbd6 BTB domain interface on the opposite side of the cullin N-terminal stalk. Thus bound, the C-terminal linker and Zn-finger region may extend back across the complex to the E2 bound to the cullin C-terminal domain for ubiquitin transfer onto a yet to be identified lysine residue. Alternatively, encounters between Cul3/Btbd6/Plzf trimers in the cytoplasm might result in the re-association of Btbd6 homodimers accompanied by a ‘hand-off’ of the displaced Plzf to the ubiquitintransfer site.

Taken together, our biochemical and modeling data, along with the demonstration of a dual role for Btbd6 as a regulator of nuclear import now suggest an integrated model for developmental stage-specific down-regulation of transcriptional repression by Plzf (Figure 4C). Here, iterative cycles of Btbd6 nuclear entry, BTB-domain exchange with Plzf and export of the heterodimeric complexes to the cytoplasm precede docking with Cul3 and eventual ubiquitination and degradation. Protomer exchange between structurally related homodimers to form heterodimeric species is often slow and it seems likely that Plzf removal from the nuclear compartment may be rather inefficient. Nonetheless, this cyclical process is embedded in a developmentally regulated transcriptional program whereby activation of neurogenin-dependent transcription of Btbd6 results in reduced levels of nuclear Plzf through the mechanisms we have described. Down-regulation of Plzf then relieves its repression of neurogenin establishing a positive-feedback loop that ensures a quantitative and timely removal of Plzf-dependent transcriptional repression.

## Materials and Methods

### Protein Expression and Purification

Zebrafish Plzfa, Btbd6a and Cul3a deletions were cloned in pPROEX HTa vector encoding N-terminal 6xHis tag and a TEV protease cleavage site. The constructs were transformed in Escherichia coli BL21 Competent Cells and grown at 18°C overnight with 1 mM IPTG for protein expression. Proteins were purified by Ni-NTA affinity purification, ion exchange and gel filtration chromatography. Btbd6 deletions are: BTB domain residue 119-234; BTB-BACK residue 119-361; PHR residue 387-540. Plzf deletions: BTB domain residue 8-126; Plzf ΔZF residue 8-401) and Cul3 deletion NTD residue 1-379.

### Microscale thermophoresis (MST)

Microscale thermophoresis (MST) measurements were performed using a NanoTempervMonolithTM NT.115 instrument (NanoTemper Technologies GmbH, München, Germany). Protein samples were labelled with the amine reactive dye NT-647 using the Monolith™ NT.115 Protein Labeling Kit RED-NHS. Labeling levels (generally in the range 0.3-0.4 dye molecules per protein) were determined using calculated extinction coefficients and ε647 =250000 M-1cm-1 for the dye concentration. In a typical experiment 20 μl aliquots of a 100 nM stock solution of labelled protein were mixed with 20 μl aliquots of a serial dilution of binding partner. These solutions were then loaded into standard treated capillaries and MST measurements were performed at 25oC using 20-40% LED power and 40-60% IR-Laser power. The laser Laser-On time was 30 seconds and Laser-Off time 5 seconds. All measurements were performed at least five times.

### Size Exclusion Chromatography Multi-Angle Laser Light Scattering (SEC-MALLS)

Size exclusion chromatography coupled to multi-angle laser light scattering (SECMALLS) was used to determine the molar mass distributions of the BTB-Btbd6/Cul3- and BTB-Plzf/Cul3-NTD complexes. Samples (100 μl) ranging in protein concentration from 15-250 μM were applied to a Superdex 200 10/300 GL column (GE Healthcare) mounted on a Jasco HPLC equilibrated in 25 mM Tris-HCl, 150 mM NaCl and 0.5 mM TCEP, pH 8.0, at a flow rate of 0.5 mL/min. The scattered light intensity and the protein concentration of the column elute were recorded using a DAWN-HELEOS-II laser photometer and OPTILAB-TrEX differential refractometer respectively. The weight-averaged molar mass of material contained in chromatographic peaks was determined from the combined data from both detectors using the ASTRA software version 6.1.1 (Wyatt Technology Corp., Santa Barbara, CA, USA).

### Mass Spectrometry

HEK 293T cells where transfected with Btbd6 pCS2 with FLAG epitope. Cells were harvested 48 hr after transfection and treated with E1a lysis buffer. The cell lysis was immunoprecipitated with FLAG antibody (Sigma F7425, Rabbit anti FLAG). The cell lysates was incubated with protein-G beads for 4 hr, and then washed 3 times. Samples were then loaded on an SDS-PAGE gel and run for 1 cm, and whole well were extracted and treated with trypsin. Cell Culture and imaging HEK293T cells were grown in DMEM media with 10% fetal calf serum (FCS), and incubated at 37C. Btbd6 wt and mutants were cloned in pEGFP1 encoding an N-terminal GFP. Cell transfection was done using FuGENE (Promega). For studying Btbd6 LR-NES, leptomycin B (LMB) was added at a concentration 2 ng/ml simultaneously with 15 ug/ml cycloheximide, and incubated for 3hr. Cells were then fixed by 2% v/v paraformaldehyde treatment, mounted and then analysed and imaged using Zeiss ApoTome.2 microscope. Images were recorded using a 100X oil objective.

## Abbreviations

Plzf: promyelocytic zinc-finger transcription factor
MST: microscale thermophoresis
BTB: Bric - Bric a brac/Tramtrack/Broad complex
PHR: PAM/Highwire/RPM-1.

## Acknowledgements

SJS and DAW are supported by the Francis Crick Institute which receives its core funding from Cancer Research UK (FC001156 and FC001217), the UK Medical Research Council (FC001156 and FC001217), and the Wellcome Trust (FC001156 and FC001217).

## Declaration

The authors declare they have no conflicts of interest.

## Supplementary Information

**Figure S1.**
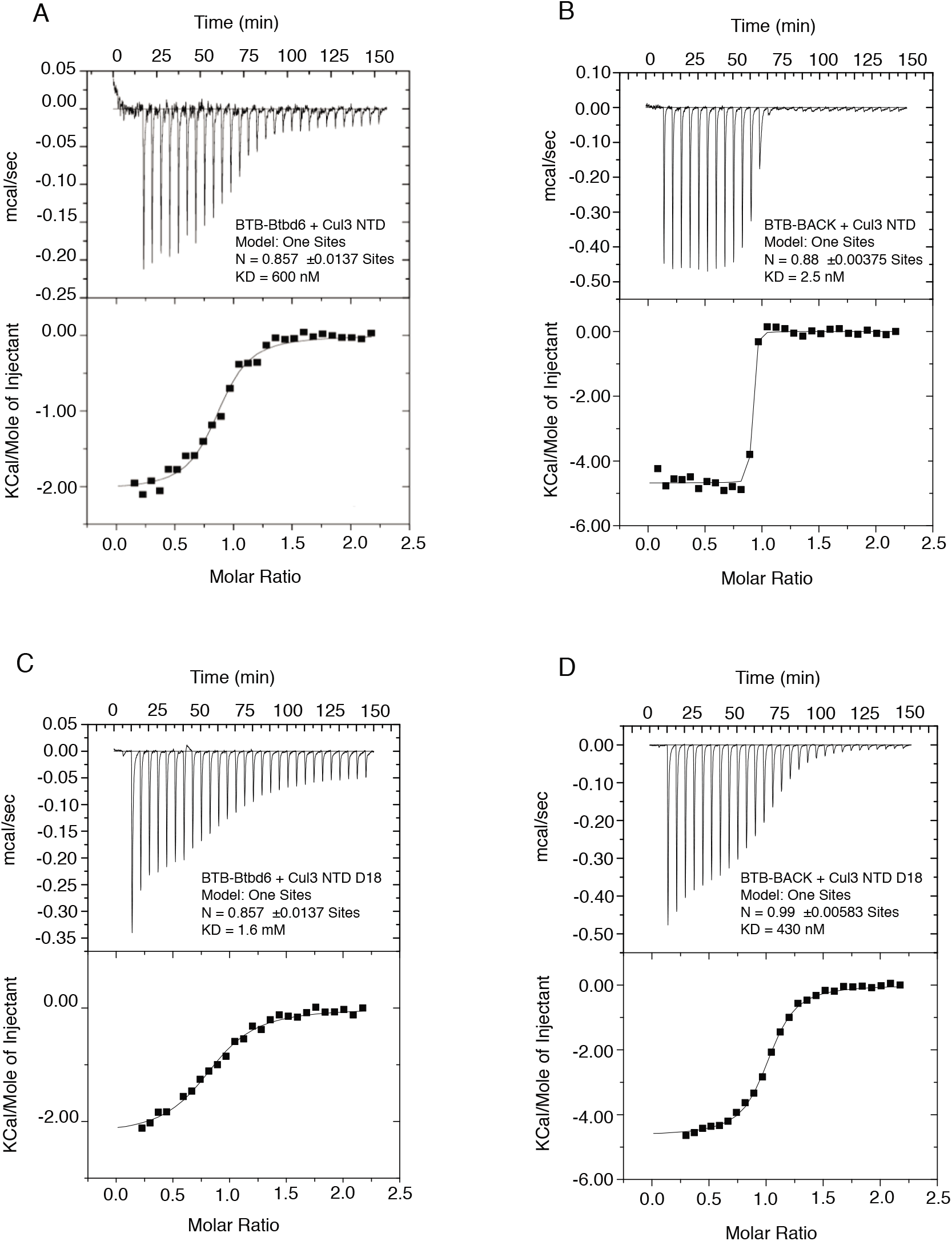
A and B) ITC analysis of BTB-Btbd6 and BTB-BACK interactions with Cul3NTD, showing that the addition of the BACK domain strongly enhances the interaction with Cul3 NTD. C and D) ITC data showing that deleting the first 18aa of Cul3 reduced the strength of the interaction with BTB-Btbd6 and BTB-BACK.

**Figure S2.**
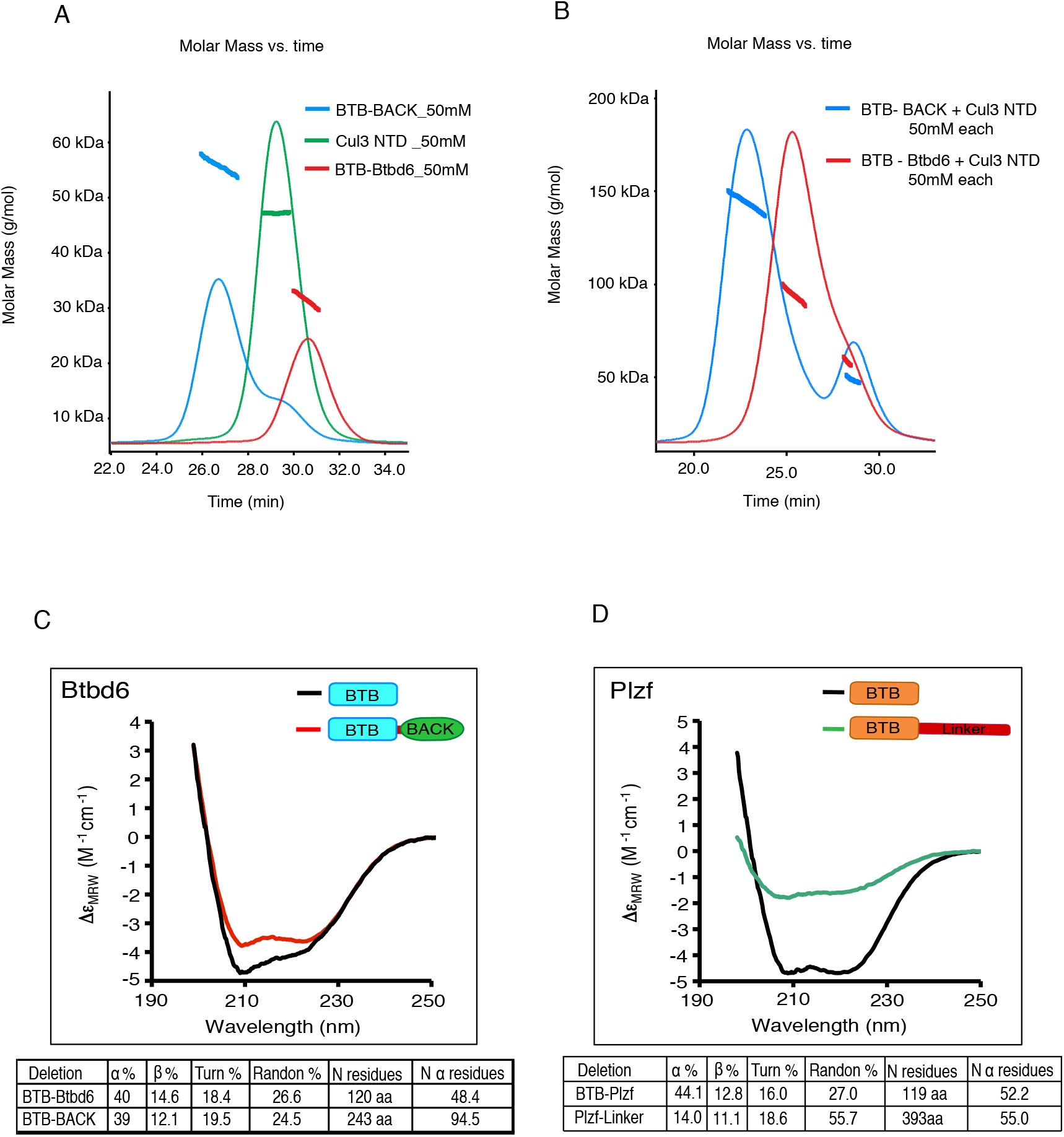
A and B) SEC-MALS experiments on BTB-Btbd6, BTB-BACK and Cul3 NTD individually (A), and in the mixtures BTB-Btbd6+Cul3 NTD or BTBBACK+Cul3 NTD (B). C) Circular dichroism on BTB-Btbd6, BTB-BACK, in comparison with BTB-Plzf (D), showing the BACK domain of BTB-Btbd6 is helical, whereas the linker following the BTB-Plzf lacks helical structure.

### Supplementary Materials & Methods

#### Isothermal Titration Calorimetry (ITC)

ITC was carried out using a VP-ITC calorimeter (MicroCal, USA). Proteins were equilibrated with a buffer containing 20 mM Tris-HCl pH 8.3, 150 mM NaCl, 0.5 mM TCEP. Typically, 10 μl aliquots of protein domains (BTB-Btbd66 or BTB-BACK) at a syringe concentration of 300 μM were titrated over 30 injections into Cul3 protein at a concentration in the ITC cell of 30 μM. Data were corrected for heats of dilution and analysed using the Origin 5.0 software.

#### Circular Dichroism (CD)

Far-UV CD spectra were recorded on a Jasco J-815 spectropolarimeter fitted with a cell holder thermostatted by a CDF-426S Peltier unit. All CD measurements were made in 25mM Tris, 100mM NaCl and 0.5mM TCEP using fused silica cuvettes with 1-mm path length (Hellma, Jena, Germany). The spectra were typically recorded with 0.1-nm resolution and baseline corrected by subtraction of the appropriate buffer spectrum. CD intensities are presented as the CD absorption coefficient calculated on a mean residue weight basis (DeMRW). Secondary structure content was estimated using methods described previously [1].

## References

[1] Pickart CM. Mechanisms underlying ubiquitination. Annu Rev Biochem. 2001;70:503–33.

[2] Michel MA, Swatek KN, Hospenthal MK, Komander D. Ubiquitin Linkage-Specific Affimers Reveal Insights into K6-Linked Ubiquitin Signaling. Mol Cell. 2017;68:233–46 e5).

[3] Deshaies RJ. SCF and Cullin/Ring H2-based ubiquitin ligases. Annu Rev Cell Dev Biol. 1999;15:435–67.

[4] Pintard L, Willems A, Peter M. Cullin-based ubiquitin ligases: Cul3-BTB complexes join the family. EMBO J. 2004;23:1681–7.

[5] Brown NG, VanderLinden R, Watson ER, Weissmann F, Ordureau A, Wu KP, et al. Dual RING E3 Architectures Regulate Multiubiquitination and Ubiquitin Chain Elongation by APC/C. Cell. 2016;165:1440–53.

[6] Dubiel W, Dubiel D, Wolf DA, Naumann M. Cullin 3-Based Ubiquitin Ligases as Master Regulators of Mammalian Cell Differentiation. Trends Biochem Sci. 2018;43:95–107.

[7] Zheng N, Schulman BA, Song L, Miller JJ, Jeffrey PD, Wang P, et al. Structure of the Cul1-Rbx1-Skp1-F boxSkp2 SCF ubiquitin ligase complex. Nature. 2002;416:703–9.

[8] Xu L, Wei Y, Reboul J, Vaglio P, Shin TH, Vidal M, et al. BTB proteins are substrate-specific adaptors in an SCF-like modular ubiquitin ligase containing CUL-3. Nature. 2003;425:316–21.

[9] Zhuang M, Calabrese MF, Liu J, Waddell MB, Nourse A, Hammel M, et al. Structures of SPOP-substrate complexes: insights into molecular architectures of BTBCul3 ubiquitin ligases. Mol Cell. 2009;36:39–50.

[10] Tong KI, Katoh Y, Kusunoki H, Itoh K, Tanaka T, Yamamoto M. Keap1 recruits Neh2 through binding to ETGE and DLG motifs: characterization of the two-site molecular recognition model. Mol Cell Biol. 2006;26:2887–900.

[11] La M, Kim K, Park J, Won J, Lee JH, Fu YM, et al. Daxx-mediated transcriptional repression of MMP1 gene is reversed by SPOP. Biochem Biophys Res Commun. 2004;320:760–5.

[12] Errington WJ, Khan MQ, Bueler SA, Rubinstein JL, Chakrabartty A, Prive GG. Adaptor Protein Self-Assembly Drives the Control of a Cullin-RING Ubiquitin Ligase. Structure. 2012.

[13] Canning P, Cooper CD, Krojer T, Murray JW, Pike AC, Chaikuad A, et al. Structural basis for Cul3 protein assembly with the BTB-Kelch family of E3 ubiquitin ligases. J Biol Chem. 2013;288:7803–14.

[14] van Geersdaele LK, Stead MA, Harrison CM, Carr SB, Close HJ, Rosbrook GO, et al. Structural basis of high-order oligomerization of the cullin-3 adaptor SPOP. Acta Crystallogr D Biol Crystallogr. 2013;69:1677–84.

[15] Sobieszczuk DF, Poliakov A, Xu Q, Wilkinson DG. A feedback loop mediated by degradation of an inhibitor is required to initiate neuronal differentiation. Genes Dev. 2010;24:206–18.

[16] Bury FJ, Moers V, Yan J, Souopgui J, Quan XJ, De Geest N, et al. Xenopus Btbd6and its Drosophila homologue lute are required for neuronal development. Dev Dyn. 2008;237:3352–60.

[17] Stogios PJ, Downs GS, Jauhal JJ, Nandra SK, Prive GG. Sequence and structural analysis of BTB domain proteins. Genome biology. 2005;6:R82.

[18] Pisani DF, Coldefy AS, Elabd C, Cabane C, Salles J, Le Cunff M, et al. Involvement of BTBD1 in mesenchymal differentiation. Experimental cell research. 2007;313:2417–26.

[19] Xu L, Yang L, Moitra PK, Hashimoto K, Rallabhandi P, Kaul S, et al. BTBD1 and BTBD2 colocalize to cytoplasmic bodies with the RBCC/tripartite motif protein, TRIM5delta. Experimental cell research. 2003;288:84–93.

[20] Siggs OM, Beutler B. The BTB-ZF transcription factors. Cell Cycle. 2012;11:3358–69.

[21] Beaulieu AM, Sant’Angelo DB. The BTB-ZF family of transcription factors: key regulators of lineage commitment and effector function development in the immune system. J Immunol. 2011;187:2841–7.

[22] Cullinan SB, Gordan JD, Jin J, Harper JW, Diehl JA. The Keap1-BTB protein is an adaptor that bridges Nrf2 to a Cul3-based E3 ligase: oxidative stress sensing by a Cul3-Keap1 ligase. Mol Cell Biol. 2004;24:8477–86.

[23] Furukawa M, He YJ, Borchers C, Xiong Y. Targeting of protein ubiquitination by BTB-Cullin 3-Roc1 ubiquitin ligases. Nat Cell Biol. 2003;5:1001–7.

[24] Mathew R, Seiler MP, Scanlon ST, Mao AP, Constantinides MG, Bertozzi-Villa C, et al. BTB-ZF factors recruit the E3 ligase cullin 3 to regulate lymphoid effector programs. Nature. 2012.

[25] la Cour T, Kiemer L, Molgaard A, Gupta R, Skriver K, Brunak S. Analysis and prediction of leucine-rich nuclear export signals. Protein Eng Des Sel. 2004;17:527–36.

[26] Bogerd HP, Fridell RA, Benson RE, Hua J, Cullen BR. Protein sequence requirements for function of the human T-cell leukemia virus type 1 Rex nuclear export signal delineated by a novel in vivo randomization-selection assay. Mol Cell Biol. 1996;16:4207–14.

[27] Kigoshi Y, Tsuruta F, Chiba T. Ubiquitin ligase activity of Cul3-KLHL7 protein is attenuated by autosomal dominant retinitis pigmentosa causative mutation. J Biol Chem. 2011;286:33613–21.

[28] Dong X, Biswas A, Suel KE, Jackson LK, Martinez R, Gu H, et al. Structural basis for leucine-rich nuclear export signal recognition by CRM1. Nature. 2009;458:1136–41.

[29] Haasen D, Kohler C, Neuhaus G, Merkle T. Nuclear export of proteins in plants: AtXPO1 is the export receptor for leucine-rich nuclear export signals in Arabidopsis thaliana. Plant J. 1999;20:695–705.

[30] Neville M, Stutz F, Lee L, Davis LI, Rosbash M. The importin-beta family member Crm1p bridges the interaction between Rev and the nuclear pore complex during nuclear export. Curr Biol. 1997;7:767–75.

[31] Fukuda M, Asano S, Nakamura T, Adachi M, Yoshida M, Yanagida M, et al. CRM1 is responsible for intracellular transport mediated by the nuclear export signal. Nature. 1997;390:308–11.

[32] Fornerod M, Ohno M, Yoshida M, Mattaj IW. CRM1 is an export receptor for leucine-rich nuclear export signals. Cell. 1997;90:1051–60.

[33] Stead MA, Wright SC. Structures of heterodimeric POZ domains of Miz1/BCL6 and Miz1/NAC1. Acta Crystallogr F Struct Biol Commun. 2014;70:1591-

## Supplementary References

[1] Sreerama, N., and Woody, R.W. (2000). Estimation of protein secondary structure from circular dichroism spectra: Comparison of CONTIN, SELCON, and CDSSTR methods with an expanded reference set. Anal. Biochem. 287: 252–260.

