## Supplemental figures for "Btbd6-dependent Plzf recruitment to Cul3 E3 ligase complexes through BTB domain heterodimerization"

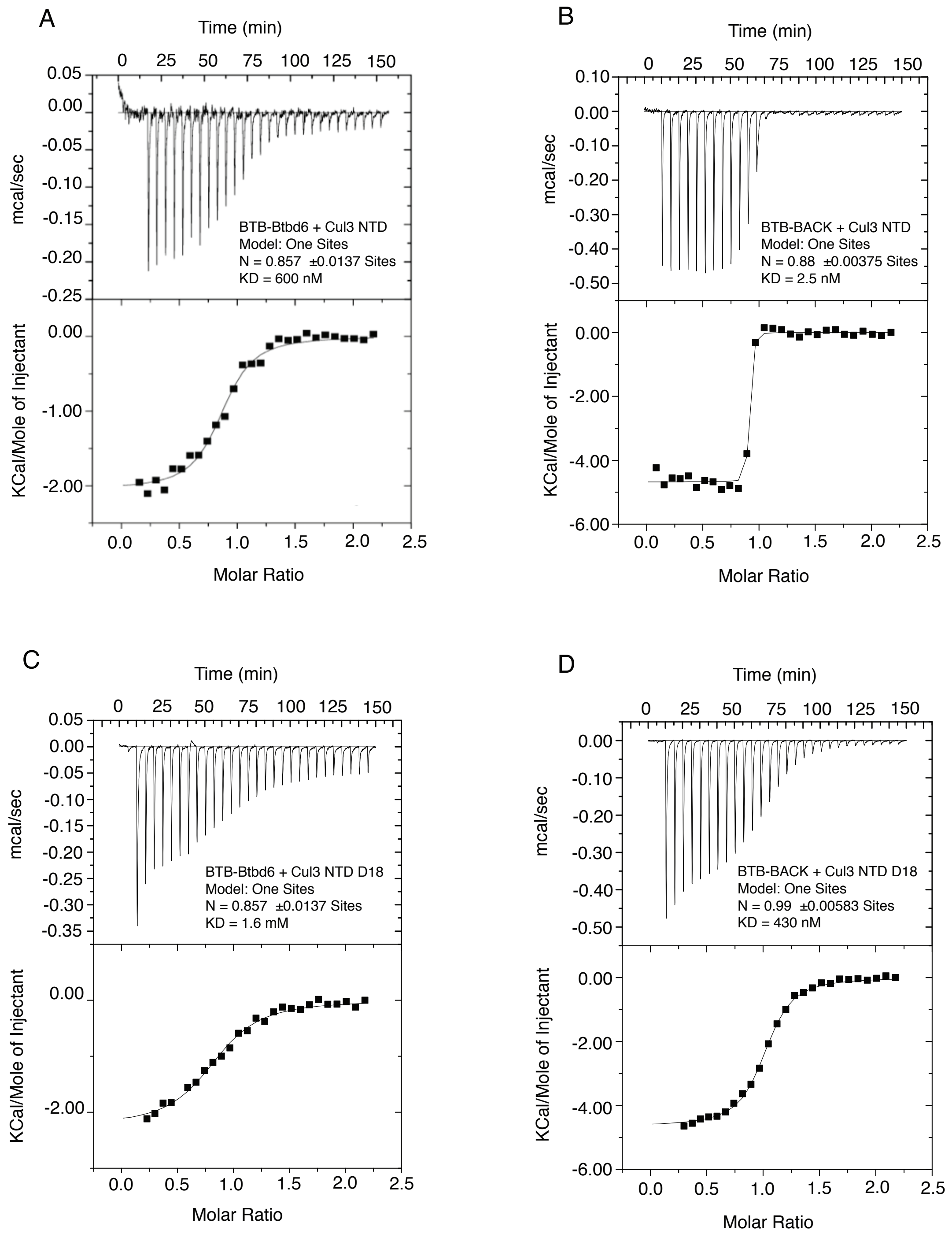


**Figure S1 -** A and B) ITC analysis of BTB-Btbd6 and BTB-BACK interactions with Cul3NTD, showing that the addition of the BACK domain strongly enhances the interaction with Cul3 NTD. C and D) ITC data showing that deleting the first 18aa of Cul3 reduced the strength of the interaction with BTB-Btbd6 and BTB-BACK.

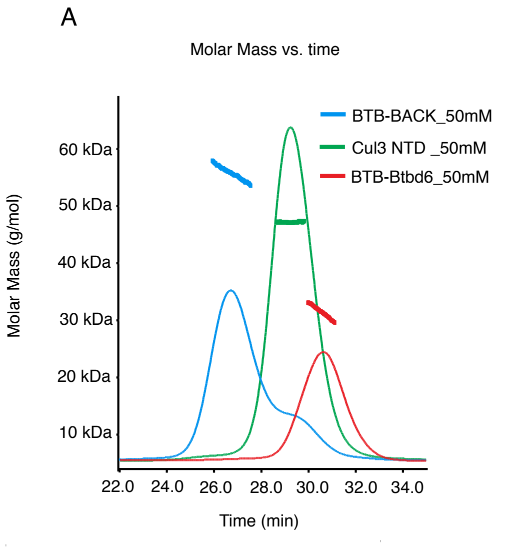


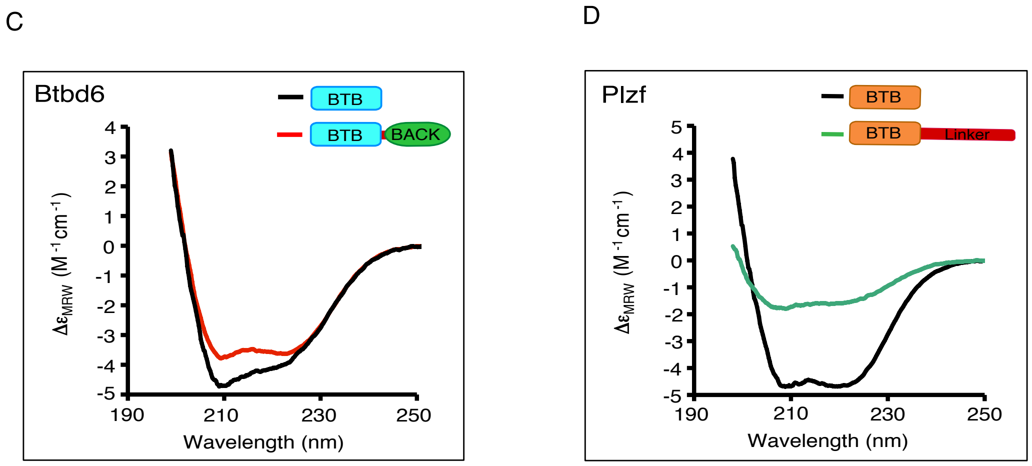


| Deletion | α % | β % | Turn % | Randon % | N residues | N α residues |
|---|---|---|---|---|---|---|
| BTB-Btbd6 | 40 | 14.6 | 18.4 | 26.6 | 120 aa | 48.4 |
| BTB-BACK | 39 | 12.1 | 19.5 | 24.5 | 243 aa | 94.5 |

| Deletion | α % | β % | Turn % | Randon % | N residues | N α residues |
|---|---|---|---|---|---|---|
| BTB-Plzf | 44.1 | 12.8 | 16.0 | 27.0 | 119 aa | 52.2 |
| Plzf-Linker | 14.0 | 11.1 | 18.6 | 55.7 | 393aa | 55.0 |

**Figure S2 -** A and B) SEC-MALS experiments on BTB-Btbd6, BTB-BACK and Cul3 NTD individually (A), and in the mixtures BTB-Btbd6+Cul3 NTD or BTBBACK+Cul3 NTD (B). C) Circular dichroism on BTB-Btbd6, BTB-BACK, in comparison with BTB-Plzf (D), showing the BACK domain of BTB-Btbd6 is helical, whereas the linker following the BTB-Plzf lacks helical structure.

### Supplementary Materials & Methods

#### Isothermal Titration Calorimetry (ITC)

ITC was carried out using a VP-ITC calorimeter (MicroCal, USA). Proteins were equilibrated with a buffer containing 20 mM Tris-HCl pH 8.3, 150 mM NaCl, 0.5 mM TCEP. Typically, 10 μl aliquots of protein domains (BTB-Btbd66 or BTB-BACK) at a syringe concentration of 300 μM were titrated over 30 injections into Cul3 protein at a concentration in the ITC cell of 30 μM. Data were corrected for heats of dilution and analysed using the Origin 5.0 software.

#### Circular Dichroism (CD)

Far-UV CD spectra were recorded on a Jasco J-815 spectropolarimeter fitted with a cell holder thermostatted by a CDF-426S Peltier unit. All CD measurements were made in 25mM Tris, 100mM NaCl and 0.5mM TCEP using fused silica cuvettes with 1-mm path length (Hellma, Jena, Germany). The spectra were typically recorded with 0.1-nm resolution and baseline corrected by subtraction of the appropriate buffer spectrum. CD intensities are presented as the CD absorption coefficient calculated on a mean residue weight basis (DeMRW). Secondary structure content was estimated using methods described previously [1].

#### Supplementary References

[1] Sreerama, N., and Woody, R.W. (2000). Estimation of protein secondary structure from circular dichroism spectra: Comparison of CONTIN, SELCON, and CDSSTR methods with an expanded reference set. Anal. Biochem. 287: 252-260.
